# New insights from 33,813 publicly available metagenome-assembled-genomes (MAGs) assembled from the rumen microbiome

**DOI:** 10.1101/2021.04.02.438222

**Authors:** Mick Watson

## Abstract

Recent rumen microbiome metagenomics papers^1,2^ have published hundreds of metagenome-assembled genomes (MAGs), comparing them to 4,941 MAGs published by Stewart *et al*^3^ in order to define novelty. However, there are many more publicly available MAGs from ruminants. In this paper, for the first time, all available resources are combined, catalogued and de-replicated to define putative species-level bins. As well as providing new insights into the constitution of the rumen microbiome, including an updated estimate of the number of microbial species in the rumen, this work demonstrates that a lack of community-adopted standards for the release and annotation of MAGs hinders progress in microbial ecology and metagenomics.

## Background

Recent advances in metagenomic assembly and binning have given rise to very large collections of metagenome-assembled genomes (MAGs), which often represent the only genomic information about a microbial strain or species that has not yet been cultured. These MAGs provide essential insight into the functional potential of individual strains and species, as well as the microbiomes they inhabit.

As globally important food-producing animals, ruminants are the subject of intense research, particularly as the microbiome in the rumen is primarily responsible for the breakdown of recalcitrant plant material into nutrients that the host can absorb. Peng *et al*^1^ and Gharechahi *et al*^2^ both recently published hundreds of rumen-derived MAGs, and compared them to 4941 rumen MAGs from Stewart *et al*^3^, and a previously published culture collection^4^. However, many more rumen-derived MAGs are publicly available, including 15 MAGs released by Hess *et al*^5^, 79 from Solden *et al*^6^, 99 from Svartstrom *et al*^7^, 251 from Parks *et al*^8^ (which analysed rumen data from Wallace *et al*^9^ and Shi *et al*^10^), 1200 from Wilkinson *et al*^11^, 4199 released by Glendinning *et al*^12^ (391 of which were the subject of the publication), and 20469 released by Stewart *et al*^3^ (4941 of which were the subject of the publication).

The 719 MAGs from Peng *et al* and the 538 MAGs from Gharechahi *et al* were compared against the full set of existing 32,557 rumen MAGs that were publicly available at the time of publication, in addition to 460 cultured genomes from the Hungate collection. Using permissive settings designed to reduce the number of false-positive species-level bins, the entire set of rumen microbial genomes was de-replicated to produce a set of putative microbial species-level bins. In addition to providing new taxonomic insights into the rumen microbiome, this work also demonstrates that the lack of a single, community-adopted repository for metagenomic bins and MAGs; the use of non-INSDC data repositories; a lack of community-adopted standards for MAG annotation; and a lack of enforcement of standards by journals, are all barriers to future research in metagenomics.

## Methods

### Data collection

MAGs from Peng *et al*, Gharechahi *et al,* Stewart *et al,* Stewart *et al*, Wilkinson *et al*, Parks *et al* and Glendinning e*t al* were downloaded from the ENA or NCBI SRA databases using BioProject identifiers listed in the relevant publication. 460 isolate genomes were downloaded from Seshadri *et al* (those which were present in INSDC databases at the time of download). MAGs from Hess *et al* were downloaded from the NERSC portal^13^. MAGs from Solden *et al* were provided by the authors via Google Drive. MAGs from Svartstrom *et al* were created by parsing the FASTA headers of metagenome assemblies released under the BioProject accession listed in the publication.

Stewart *et al* and Glendinning *et al* are noteworthy as they released large numbers of assemmbly bins (20,469 and 4,199 respectively) that were not described as part of the paper, instead choosing to select high quality MAGs from those bins for further analysis and description. However, those datasets are still useful and were included in the analysis.

Completeness and contamination scores for Stewart *et al,* Stewart *et al*, and Glendinning *et al* were retrieved from the ENA using the BioProject identifiers listed in the relevant publication. Completeness and contamination scores for Peng *et al* and Gharechahi *et al* were retrieved from the publication and linked to the published sequences using assembly statistics. Completeness and contamination scores for Seshadri *et al*, Solden *et al*, Svartstrom *et al*, Hess *et al* and Parks *et al* were calculated using CheckM^14^. Finally, completeness and contamination scores for Wilkinson *et al* were retrieved from the authors.

### Dereplication

The software dRep^15^ was used to de-replicate 33,813 publicly available FASTA files representing MAGs and isolate genomes from ruminants. The parameters used were:

- --S_algorithm fastANI
- --multiround_primary_clustering
- -comp 50
- -con 10
- -sa 0.95
- -nc 0.3

These settings delineate species-level bins if genomes have lower than 95% ANI across 30% of their length. The low length parameter (30%) recognizes that incomplete genomes may not overlap along a large proportion of their shared sequence, and using a higher value may split genomes which belong to the same species and over-estimate the number of species. A 30% coverage threshold has been used in previous large-scale MAG studies which involved 50% complete genomes^16^.

### Taxonomic classification and phylogeny

MAGpy^17^ and GTDB-Tk^18^ (with the “classify_wf” option) were used to assign a taxonomy to the “winning” MAGs from dRep. To create a representative phylogenetic tree, genomes that were at least 90% complete and less than 5% contaminated were used as input to PhyloPhlAn^19^. Proteins were predicted for the genomes with Prodigal^20^ and PhyloPhlAn was run with options:

- -d phylophlan
- -t a
- -f supermatrix_aa.cfg
- --diversity low
- --fast
- --verbose
- --maas phylophlan_substitution_models/phylophlan.tsv
- --remove_fragmentary_entries
- --fragmentary_threshold 0.85

A plot of the phylogenetic tree was created using GraphlAn^21^ with genomes coloured by the Phylum assigned by GTDB-Tk.

## Results

After de-replication using the parameters above, there remained 7,533 putative species-level bins. Treating all data released by a single publication as a single dataset (n=10), 4,794 were singletons, representing putative microbial species that are represented in only one dataset. Conversely, 2,739 putative microbial species were seen in more than one dataset. There were no species-level bins present in all datasets. The maximum number of datasets any species-level bins were present in was 7, and only three bins were present in these 7 datasets: two from Stewart *et al*, and one from Gharechahi *et al*. Results of the de-replication process are summarized in supplementary data 1.

Of the “winners”, 4482 came from Stewart *et al*, 728 from Wilkinson *et al*, 694 from Peng *et al*, 660 from Glendinning *et al*, 358 from Gharechahi *et al*, 313 from Seshadri *et al*, 149 from Parks *et al*, 78 from Svartstrom *et al*, 62 from Solden *et al* and 9 from Hess *et al.*

Using the Chao 1 index, we update the estimate of the number of microbial species in the rumen (previously estimated by Stewart *et al*^3^) to 13,616.

Of the 7,533 species-level bins, GTDB-Tk identified 155 as *Archaea*. The majority of these were assigned to the Phylum *Methanobacteriota* (119; 77%), with 31 being assigned to *Thermoplasmatota* and 5 to *Halobacteriota*. GTDB-Tk was unable to assign a Genus to eight (5%) of the Archaeal MAGs, and was unable to assign a species to 110 (71%). GTDB-Tk assigned 7,378 species-level bins to the *Bacteria* domain. These spanned 28 different Phyla, the most popular being *Firmicutes_A* (3339; 45%), *Bacteroidota* (1671; 23%), *Firmicutes* (807; 11%), *Proteobacteria* (299; 4%) and *Verrucomicrobiota* (248; 3%). Of particular interest as high fibre-degrading microbes, the dataset also contains 63 members assigned to the phylum *Fibrobacterota*. Of the 7,378 Bacterial species-level bins, GTDB-Tk was unable to assign a Class to one, unable to assign an Order to 11, unable to assign a Family to 99, unable to assign a Genus to 1,087 (15%) and unable to assign a Species to 5,796 (78%). The full results of the GTDB-Tk taxonomic assignments can be found in supplementary data 2.

A phylogenetic tree of the 2,696 highly complete MAGs can be seen in Figure 1. The tree is dominated by large clades of *Firmicutes_A* and *Bacteroidota*, though other significant clades exist with the MAGs spread across 26 different phyla.

**Figure 1.**
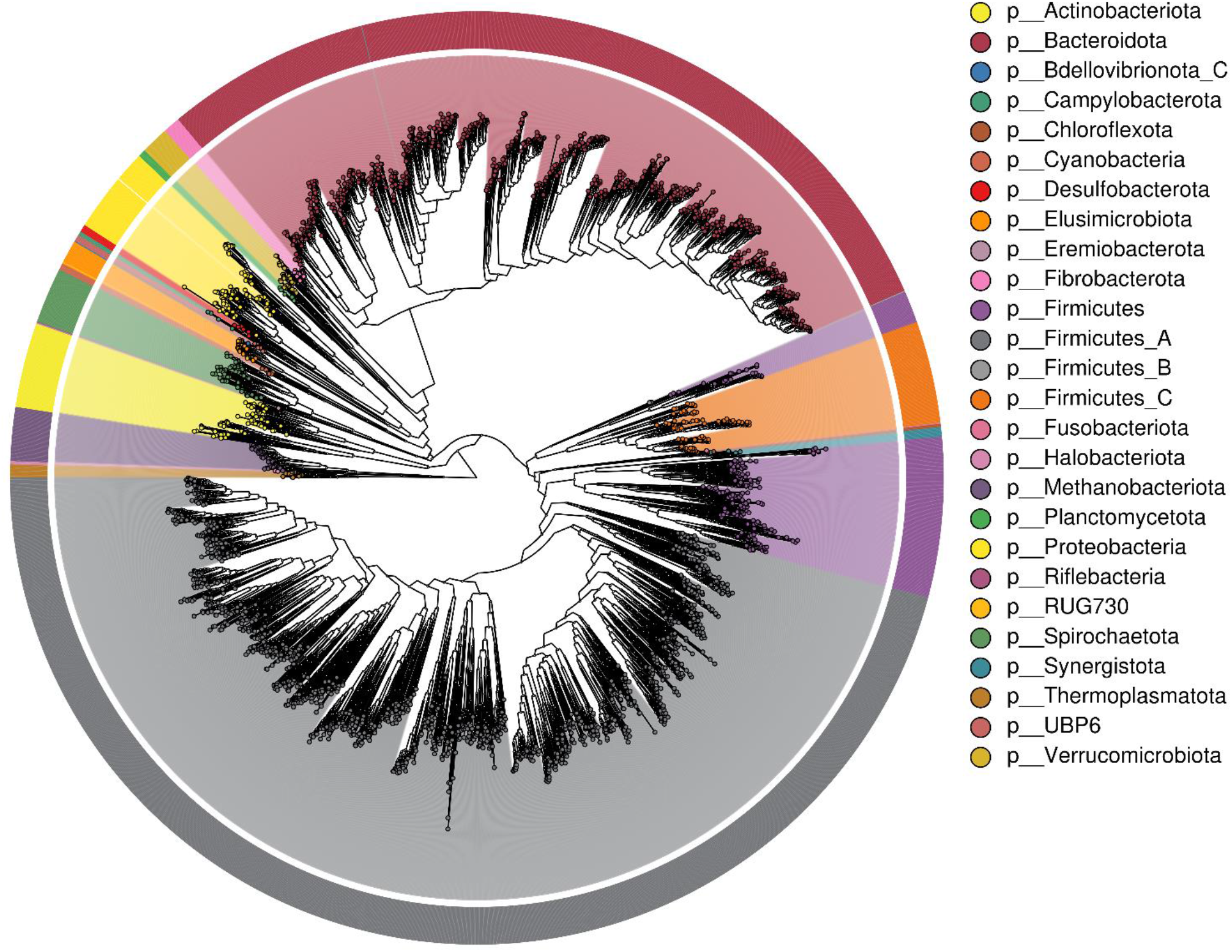
Phylogenetic tree of 2,696 species level bins chosen for high completeness (>90%) and low contamination (<5%). The tree was created using PhyloPhlAn and drawn using GraPhlAn, with taxonomic assignments from GTDB-Tk.

## Conclusions and discussion

Assembly of genomes from metagenomic sequencing is becoming routine, with new MAG datasets published frequently. In order for comparisons to be made between new and existing MAG catalogues, it should be easy to find and retrieve all MAGs from a particular biome, alongside metadata in a standardised format. The MIMAG standard attempts to define standards for metadata around MAGs^22^, but these have not been universally adopted, and cannot be applied retrospectively to historical datasets. MAGs should be deposited in INSDC databases alongside metadata adhering to the MIMAG standard, and with completeness and contamination estimates as a minimum. In addition, it would be beneficial to submit all of the following: the raw reads; the raw metagenome assemblies; all metagenome bins created during the binning process; the final set of metagenome-assembled genomes. The European Nucleotide archive (ENA) provides a suitable repository and guidance for submitting all of these^23–25^; and have metadata checklists for both binned genomes^26^ and metagenome-assembled genomes^27^ that allow for the storage of essential metadata needed to interpret and compare metagenomics data. EBI’s MGnify^28^ also provides added value services on these datasets.

There are approximately 4 billion food-producing ruminants on the planet at any one time (FAOSTAT), and as such, the rumen and its microbiome are a priority area for research globally. Recent advances in metagenomics mean we can now study the structure and function of the rumen microbiome in unprecedented detail. Here, for the first time, all data resources representing binned metagenomes from ruminants were combined and de-replicated to produce 7,533 putative species-level bins. However, combining and de-replicating partial and contaminated genomes is an inexact science, and it is difficult to effectively delineate species given the amount of missing information. Relatively permissive parameters were chosen so as to avoid over-inflation of the number of species-level bins. However it is still possible that this is an over-estimate due to the inclusion of incomplete genomes.

The fact that the majority of the species-level bins for ruminants are singletons suggests that we are yet to sample the entire species-level sequencing space of global ruminants, and the lack of a species-level core microbiota suggests a large amount of variation in the constitution of the rumen microbiome. We update the estimate of the total number of microbial species in the rumen microbiome to 13,616 and provide taxonomic labels for all known species to date, which span 26 different microbial phyla. It is essential that researchers are funded to culture this microbial diversity, and study the role of these species in ruminant productivity and health, in climate change and sustainability, and in global food security.

## Funding

The Roslin Institute forms part of the Royal (Dick) School of Veterinary Studies, University of Edinburgh. This project was supported by the Biotechnology and Biological Sciences Research Council (BBSRC; BB/N016742/1, BB/N01720X/1), including institute strategic program and national capability awards to the Roslin Institute (BB/P013759/1, BB/P013732/1, BB/J004235/1, BB/J004243/1, BBS/E/D/30002276), the Technology Strategy Board (TS/J000108/1, TS/J000116/1), and by the Scottish government as part of the 2016–2021 commission.

## Notes

### Competing Interest Statement

The authors have declared no competing interest.

